# A conserved requirement for *Fbxo7* during male germ cell cytoplasmic remodelling

**DOI:** 10.1101/563718

**Authors:** Claudia C Rathje, Suzanne J Randle, Sara Al Rawi, Benjamin M Skinner, Emma EP Johnson, Joanne Bacon, Myrto Vlazaki, Nabeel A Affara, Peter J Ellis, Heike Laman

## Abstract

Fbxo7 is the substrate-recognition subunit of an SCF-type ubiquitin E3 ligase complex. It has physiologically important functions in regulating mitophagy, proteasome activity and the cell cycle in multiple cell types, like neurons, lymphocytes and erythrocytes. Here we show that in addition to the previously-known Parkinsonian and haematopoietic phenotypes, Fbxo7-deficient male mice are completely sterile. In these males, despite successful meiosis, nuclear elongation and eviction of histones from chromatin, the developing spermatids are phagocytosed by Sertoli cells during late spermiogenesis, as the cells undergo cytoplasmic remodelling. Surprisingly, despite the loss of all germ cells, there was no evidence of the symplast formation and cell sloughing that is typically associated with spermatid death in other mouse sterility models, suggesting that novel cell death and/or cell disposal mechanisms may be engaged in Fbxo7-deficient males. Mutation of the *Drosophila* Fbxo7 orthologue, *nutcracker* (*ntc*) was previously shown to cause sterility at a similar stage of germ cell development, indicating that the requirement for Fbxo7 is conserved. The *ntc* phenotype was attributed to proteasome mis-regulation via an interaction with the proteasome regulator, DmPI31. Our data suggest rather that in mice, the requirement for Fbxo7 is either independent of its interaction with PI31, or relates specifically to cytoplasmic proteasome activity during spermiogenesis.

## Introduction

Developing male germ cells progress through a proliferative phase (spermatogonia), a meiotic phase (spermatocytes), and a spermiogenic phase (spermatids), following which spermatozoa are released through the lumen of the seminiferous tubules into the epididymis, where they undergo further maturation and acquire motility. Defects at any of these stages lead to various forms of impaired male fertility, including low sperm count (oligozoospermia), abnormal sperm shape or size (teratozoospermia), impaired sperm motility (asthenozoospermia), abnormal nuclear chromatin compaction and poor fertilisation capacity (Hess and Renato, 2008; Hecht, 1995; Jamsai and O’Bryan, 2011; Curi et al., 2003; Rathke et al., 2014). During spermiogenesis, round haploid spermatids undergo terminal differentiation to form spermatozoa, developing specialised organelles – the acrosome and flagellum – necessary for fertility and motility, respectively. This involves a dramatic morphological transformation, including nuclear compaction via the eviction of histones and their replacement with protamines, and elimination of the bulk cytoplasmic content of the developing cells. Both these changes serve to streamline the spermatozoa and improve their hydrodynamic properties to allow rapid progressive motility within the female reproductive tract. Until late spermiogenesis, spermatids remain connected by intercellular bridges, through which cytoplasmic constituents are shared among haploid spermatids (Ventela et al., 2003; Braun et al., 1989). Cytoplasmic shedding also removes these bridges and allows the individual sperm cells to separate in a process called individuation.

The disposal of excess cytoplasmic contents, including mitochondria and other organelles, is critical to many aspects of late spermatid differentiation (Sakai and Yamashina, 1989). Key elements of this process are conserved. In mammalian spermatogenesis, cytoplasmic processes from the supporting Sertoli cells invade the spermatid cytoplasm during late elongation to form the “mixed body.” Concurrently, deep invaginations known as “crypts” form within the Sertoli cell cytoplasm. Active transport of the spermatids to the base of the crypts enables the development of extensive cell-cell contacts between the Sertoli and germ cells. As remodelling progresses, branches of the invading processes engulf and phagocytose portions of the spermatid cytoplasm, resulting in the loss of around 50% of spermatid cytoplasmic volume prior to spermiation. Finally, at spermiation, the spermatids are ejected from the crypts and actively transported back to the tubule lumen. There, they shed their remaining cytoplasm as residual bodies, which are then also phagocytosed by the Sertoli cells (Sakai and Yamashina, 1989; Kerr and de Kretser, 1974; Russell et al., 1989). In *Drosophila* spermatogenesis, an actin-based individualization complex slides caudally along a group of 64 interconnected spermatids, promoting their separation and the removal of most of their cytoplasm and organelles into a membrane-bound sack, the cystic bulge, eventually discarded as a waste bag – the equivalent of the mammalian residual body (Fabian and Brill, 2012).

Processing of spermatid cytoplasm in preparation for phagocytosis by the Sertoli cells includes caspase activation (Blanco-Rodriguez and Martinez-Garcia, 1999; Arama et al., 2003; Cagan, 2003) and the degradation of cellular components by specialized variants of the proteasome (Zhong and Belote, 2007; Qian et al., 2013; Bose et al., 2014). The 20S catalytic core of a proteasome is a barrel-shaped assembly, comprised of α and β subunits. Three of the β subunits, β1, β2 and β5, have peptidase activity, while access of substrates into the core is controlled by α subunits, which recruit proteasome activators (PAs). The constitutively expressed proteasome consists of a regulatory 19S ‘lid’ associated with a 20S core particle (Voges et al., 1999; Bochtler et al., 1999), but in particular cell types or under stress conditions, alternate proteasome configurations come into play (Kniepert and Groettrup, 2014). During spermatogenesis, an alternative α4-type proteasome subunit, α4s/PSMA8, a testis-specific subunit, replaces its 20S counterpart, which is thought to enable the recruitment of an alternate lid, PA200. PA200-capped proteasomes promote the degradation of acetylated histones, which enables their removal from DNA and replacement with protamines, for enhanced nuclear compaction into the spermatid head (Qian et al., 2013; Gaucher et al., 2010). A second alternate proteasome, known as the immunoproteasome, is also expressed during spermiogenesis. It substitutes β-subunits, β1i, β2i and β5i, and has a different regulatory 11S lid (Qian *et al.*, 2013). As sperm differentiation requires major cellular remodelling and volume reduction, these alternate proteasomes are thought to play crucial roles in fashioning this specialized cell form (Rathke et al., 2014; Zhong and Belote, 2007; Kniepert and Groettrup, 2014).

In *Drosophila*, Nutcracker (*ntc*) protein is essential for spermatid individuation, and homozygous *ntc* mutant male flies are sterile. Spermatids deficient for *ntc* undergo apoptosis in late spermiogenesis at the point when individuation would normally occur. The spermatid apoptosis is associated with failure to form individuation complexes, failure to activate spermiogenesis-related caspases, and reduced proteasome activity in *ntc*-deficient testes. These defects have been ascribed to an interaction between *ntc* and proteasome binding protein PI31 (Bader et al., 2011; Bader et al., 2010; Arama et al., 2007). PI31 was discovered as an *in vitro* inhibitor of proteasome activity (Zaiss et al., 2002; McCutchen-Maloney et al., 2000), which distinguished it from other proteasome regulators, like PA200, PA28, and 19S, which all activate the 20S proteasome. However, within intact cells, PI31 effects on proteasome activity remain unclear, with contrasting results being reported for the *Drosophila* and mammalian PI31 proteins. In *Drosophila*, DmPI31 activation of the 26S proteasome is essential for sperm differentiation, and DmPI31 levels are greatly reduced in *ntc* mutant testes, indicating that DmPI31 requires a stabilizing interaction with *ntc* to achieve sufficiently high expression levels (Bader et al., 2011; Bader et al., 2010; Arama et al., 2007). However, while transgenic overexpression of DmPI31 in *ntc* mutant testes promoted caspase activation in germ cells, it failed to rescue the sterility phenotype (Bader *et al.*, 2011). It is therefore unclear whether the sterility effects of *ntc* deficiency are mediated by the *ntc* / DmPI31 interaction.

A mammalian orthologue of Nutcracker is Fbxo7, although there may be functional differences between them given the inability of human Fbxo7 to rescue the sterility of *ntc* flies (Burchell *et al.*, 2013). However, like DmPI31 and Nutcracker, mammalian PI31 and Fbxo7 heterodimerize via their FP domains (Shang et al., 2015; Kirk et al., 2008). Studies in cultured mammalian cells indicate that PI31 has little effect on constitutive proteasome activity or assembly but instead acts as a selective inhibitor of the maturation of immunoproteasomes (Li et al., 2014; Zaiss et al., 1999). The lack of a pronounced effect on proteasome activity may indicate that PI31 functions in specific contexts and/or in spatially or temporally-restricted ways. The robust, conserved interactions of Fbxo7 and PI31 with each other and also with the proteasome suggest they have roles in regulating proteasome function. Fbxo7 is a multifunctional, F-box protein with distinct activities in different cell types. In human health Fbxo7 impacts on numerous pathologies, including Parkinson’s disease, cancer and anaemia (Ding et al., 2012; Ganesh et al., 2009; van der et al., 2012; Soranzo et al., 2009; Lomonosov et al., 2011; Paisan-Ruiz et al., 2010; Di Fonzo et al., 2009; Laman, 2006; Lohmann et al., 2015). At a molecular level, Fbxo7 functions as a receptor for SCF-type E3 ubiquitin ligases and also non-canonically, as a scaffolding chaperone for other regulatory proteins. Its effects are observable in NF-κB signalling, via cIAP and TRAF2 interactions (Kuiken *et al.*, 2012), and in cell cycle regulation via Cdk6 activation and p27 stabilisation (Patel et al., 2016; Randle et al., 2015; Meziane et al., 2011; Laman, 2006). Fbxo7 has also been shown to interact with and ubiquitinate proteasome subunits, like PSMA2 (Teixeira et al., 2016; Vingill et al., 2016; Fabre et al., 2015; Bousquet-Dubouch et al., 2009).

We report here that, like *ntc* flies, male mice with a reduced expression of Fbxo7 are infertile. In these mice, developing spermatids are phagocytosed en masse by the Sertoli cells, beginning at the stage when they would normally start to shed their cytoplasm. The very few sperm that escape phagocytosis and complete maturation have grossly aberrant morphology. Thus, flies and mice both exhibit a catastrophic loss of germ cells at the onset of cytoplasmic remodelling. This is a novel form of spermatogenic failure unlike any other previously-described infertile mouse model. However, Fbxo7 mutant mice show only a slight reduction in PI31 levels and normal levels of constitutive proteasome activity in spermatids, indicating that in mouse the fertility effects of Fbxo7 mutation may be independent of its effect on PI31 and the proteasome.

## Results

### Fbxo7^LacZ/LacZ^ mice are sterile due to azoospermia

In the course of our investigations into the physiological functions of mammalian Fbxo7, we generated mice that are either heterozygous or homozygous for an allele of Fbxo7 containing a *LacZ* insertion between exons 3 and 4 of Fbxo7 (Patel et al., 2016; Randle et al., 2015). This insertion severely disrupts expression of all Fbxo7 isoforms but does not completely abolish it (Randle *et al.*, 2015), and thus the phenotype(s) of the hetero- and homozygous animals can respectively be ascribed to moderate or severe under-expression of Fbxo7. In maintaining the colony of LacZ-transgenic animals, we observed that homozygous Fbxo7^LacZ/LacZ^ males never sired any offspring, while heterozygous males and all genotypes of female were able to produce litters. In heterozygous crosses, there was a mild deficit of homozygous offspring (**Figure 1A**), suggestive of a small degree of embryonic lethality in this genotype.

**Figure 1.**
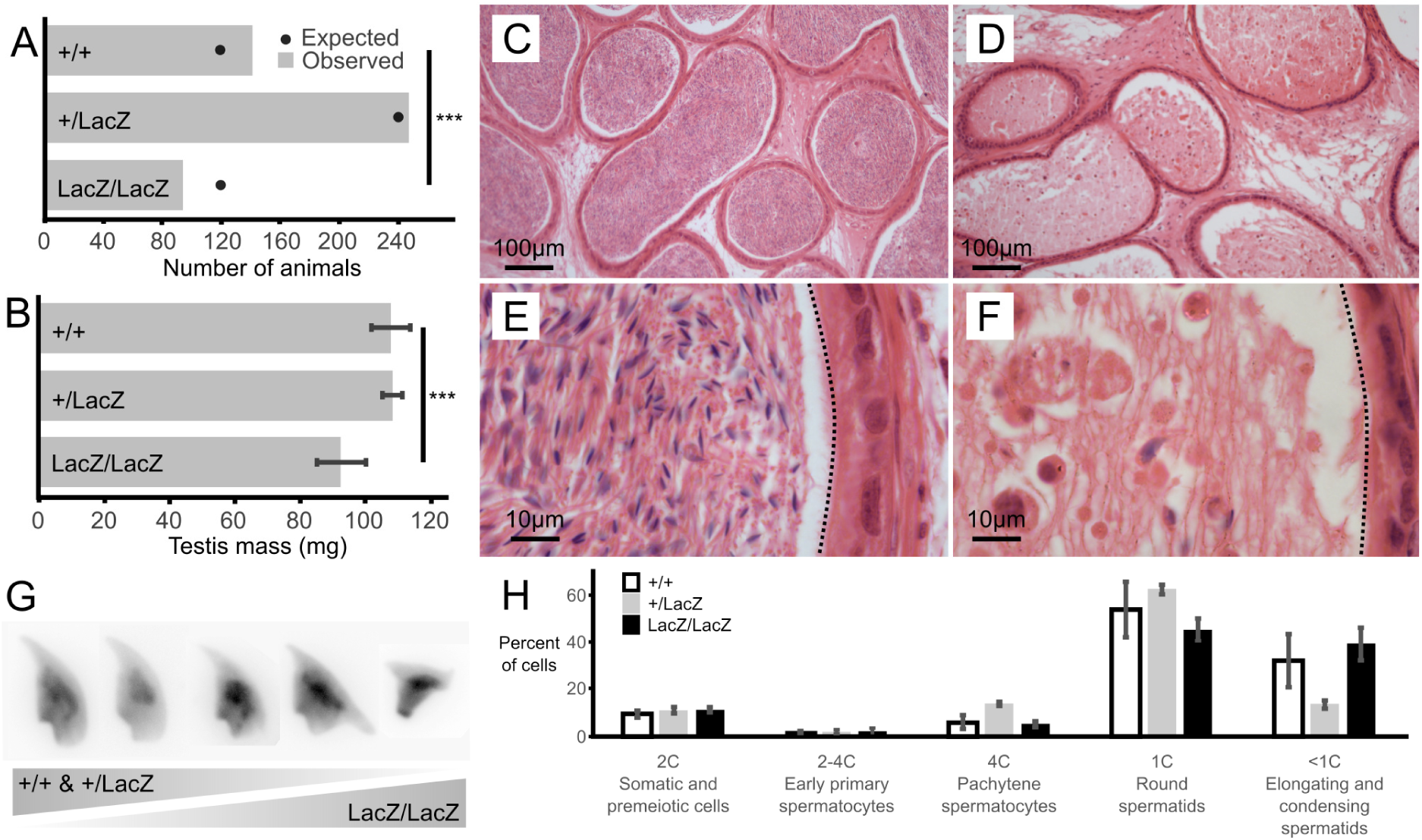
Male mice deficient for Fbxo7 are sterile. **A:** Colony data showing the genotypes of animals (n=481) born from matings between heterozygous parents. All three potential genotypes are observed in the offspring, but the proportion of homozygous Fbxo7^LacZ/LacZ^ offspring is lower than expected (*** chi-squared goodness-of fit test p= 0.007 vs Mendelian 1:2:1 expectation). **B:** Average testis weight for each genotype (n=6; WT and homozygous; n=4, heterozygous LacZ; One-way ANOVA, *** p = 0.001067)). **C:** Wild type cauda epididymis showing large numbers of stored sperm. **D:** Fbxo7^LacZ/LacZ^ cauda epididymis showing very few degenerating sperm. **E/F:** High resolution zoom of sections C/D. Dotted line indicates the border of the tubule lumen in each view. **G:** Montage of DAPI-stained sperm nuclei showing the spectrum of sperm morphologies present (see also **Supplementary Figure 1**). **H:** FACS quantitation of testis cells according to DNA content as measured by propidium iodide staining (note that highly condensed spermatids and mature sperm were not quantitated).

Initial characterisation showed a significant reduction in mean testis weight for the Fbxo7^LacZ/LacZ^ compared to heterozygous and WT males (92.4 mg vs 108.2 and 107.7 mg; **Figure 1B**), indicative of abnormal testis development. Strikingly, there were virtually no mature sperm in the lumen of the epididymis of the Fbxo7^LacZ/LacZ^ males (**Figure 1C-F**), demonstrating that these males are sterile due to azoospermia. A very few residual sperm were retrieved from dissected epididymides of Fbxo7^LacZ/LacZ^ males, with a total yield of fewer than 1,000 cells per cauda epididymis, compared to a normal count of around 10^8^ sperm cells per cauda. The residual sperm were all grossly misshapen, and a high proportion of cells showed abnormal compression of the rear of the sperm head. Heterozygous Fbxo7^LacZ/+^ males also showed a slight increase in the frequency of abnormally shaped sperm, with the most severely deformed sperm resembling the homozygous phenotype (**Figure 1G**). Using a newly-developed image analysis programme for sperm morphometry (Skinner *et al.*, 2019), we determined that the remaining sperm from Fbxo7^LacZ/LacZ^ males showed an average 14% reduction in cross-sectional area, with high morphological variability (**Supplementary Figures 1, 2**).

**Figure 2.**
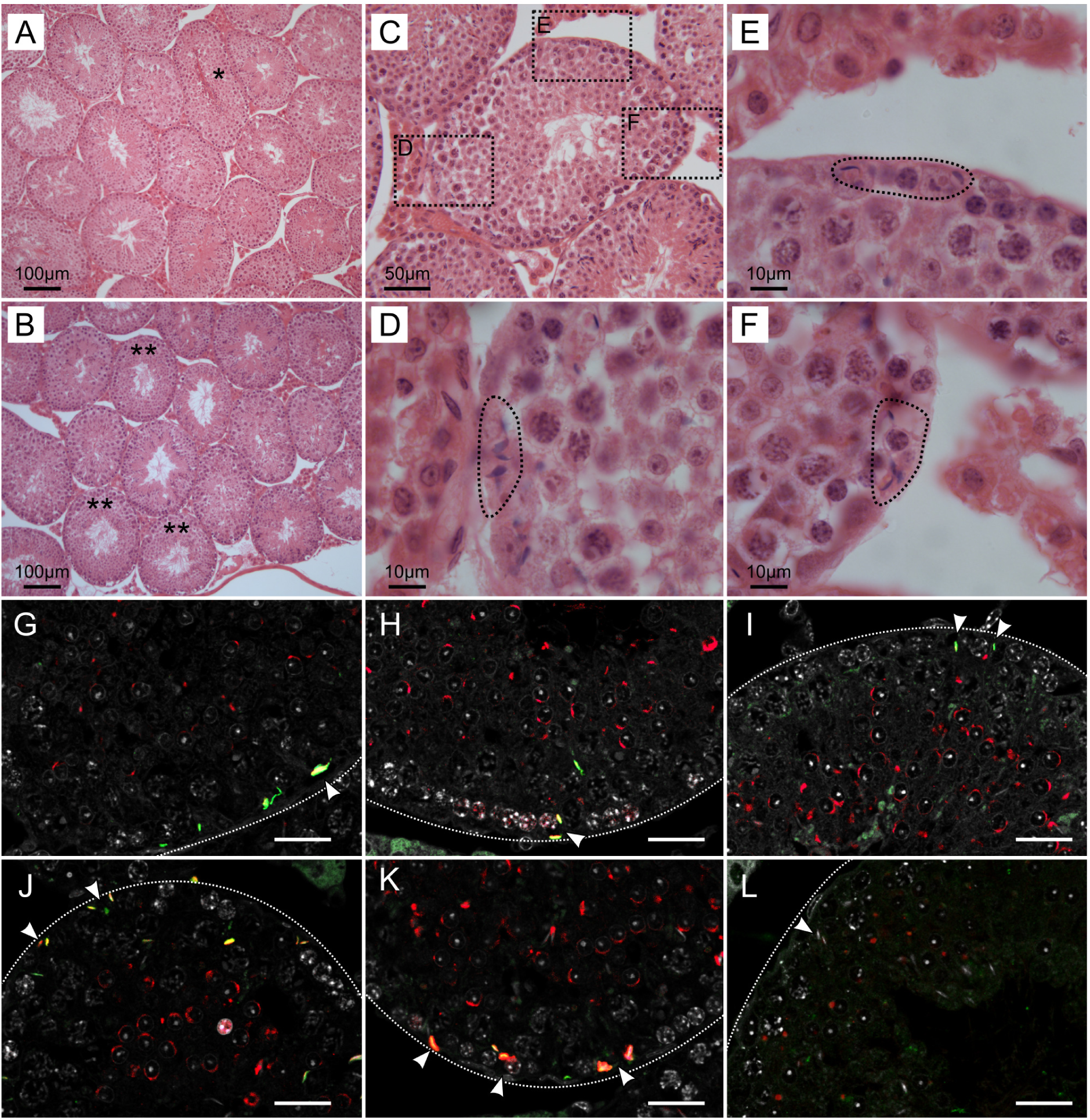
Massive loss of maturing sperm in Fbxo7 mutant males. **A-B:** Low magnification view of H&E sections from wild type **(A)** and Fbxo7^LacZ/LacZ^ testes **(B)**. * in panel **A** indicates a tubule at stage VII-VIII with sperm heads lined up at the lumen awaiting release. These were never observed in Fbxo7^LacZ/LacZ^ testes. ** in panel **B** indicates tubules with a layer of round spermatids but which lack elongating spermatids. **C:** High magnification view of a Fbxo7^LacZ/LacZ^ tubule lacking elongating spermatids: **D-F:** Close up zoom from panel C at the indicated locations. The dotted outlines indicate “graveyards” containing multiple phagocytosed spermatid heads. **G-I:** Immunofluorescent stains in Fbxo7^LacZ/LacZ^ tubules for LAMP2 (green) with PNA-lectin (red) to stage the tubules and DAPI counterstain (grey). Phagocytosed cells marked by LAMP2 were visible at tubule stage VI as indicated by the extent of the lectin-stained acrosomal cap. **J-L:** Immunofluorescent stains in Fbxo7^LacZ/LacZ^ tubules for CASP2 (green) with PNA-lectin (red) to stage the tubules and DAPI counterstain (grey). Apoptotic cells marked by CASP2 were visible at tubule stage VI as indicated by the extent of the lectin-stained acrosomal cap. At tubule stage IV (L), occasional mis-localised elongating spermatids were seen next to the basement membrane. These cells were not marked with CASP2 at this stage. For quantitation of spermatid mis-localisation, see **Figure 3**.

### Developing Fbxo7^LacZ/LacZ^ spermatids are lost during late spermiogenesis

The relatively small magnitude of the change in testis weight suggested that any germ cell abnormality was likely to only affect later stages of germ cell development. To characterise the stage of germ cell loss in Fbxo7^LacZ/LacZ^ males, we carried out an initial assessment by FACS to see if there was any gross defect in meiotic progression (**Figure 1H**). There was no significant alteration in the ratio of cells containing 1C, 2C or 4C DNA content, respectively representing haploid round spermatids, spermatogonia and primary spermatocytes, indicating no cell loss prior to the onset of spermatid elongation. Elongating and condensing spermatids/spermatozoa often appear as lower than 1C DNA content in FACS staining, but cannot be reliably quantified due to their high variability in staining parameters (Simard *et al.*, 2015).

Histological examination of adult testes using hematoxylin and eosin staining (H&E) showed limited gross changes in testis structure (**Figure 2**, **Supplementary Figure 2**). In both Fbxo7^LacZ/LacZ^ males and wild type (WT) males, pre-meiotic, meiotic and post-meiotic stages of germ cell development were all present in the testis parenchyma (**Figure 2 A,B**). However, in Fbxo7^LacZ/LacZ^ testes, very few tubules showed sperm heads adjacent to the lumen (**Figure 2 B,C**), suggesting that germ cells are lost prior to spermiation. Instead, testes from these males contained tubules with no (or virtually no) elongating cells, but where the first layer of spermatids were still round. These are tubules in the first half of the seminiferous cycle but where the late elongating cells have been lost. In these tubules, sperm heads were observed lying deep within the Sertoli cell cytoplasm near to the basement membrane, often in quite dramatic “graveyards” containing multiple cells in advanced stages of karyolysis (**Figure 2D-F**). Strikingly, however, we did not observe any formation of multinucleate symplasts or any sloughing of degenerating cells into the lumen (note also the lack of sloughed cells in the epididiymis in **Figure 1F**).

### “Graveyards” of phagocytosed Fbxo7^LacZ/LacZ^ condensing spermatids at tubule stage VI are positive for caspase-2

Since the normal fate of arrested germ cells is apoptosis followed by either phagocytosis or cell sloughing, we used fluorescent immunohistochemical staining for caspase-2 and LAMP2 (Lysosome-associated membrane protein 2) to trace these processes. Caspase 2 is an apical caspase implicated in stress-mediated germ cell death (Johnson et al., 2008; Lysiak et al., 2007; Zheng et al., 2006), while LAMP2 labels late stage phagolysosomes. In this experiment, we also used fluorescently labelled peanut agglutinin (PNA) to label the acrosomes, allowing for more detailed tubule staging (**Figure 2 G-K**). This showed that the cells in the “graveyards” were most prominent at tubule stage VI, and were positive for both LAMP2 and caspase-2. Lower-level caspase-2 staining was sometimes visible at this stage in spermatids located further from the basement membrane. We hypothesize these latter cells to be in the process of engulfment. The few remaining condensing spermatids near the lumen were still caspase-2 negative. Thus, the spermatids in the “graveyards” are apoptotic cells that have been phagocytosed by the Sertoli cells. Immunohistochemical staining for caspase-3 gave negative results in both genotypes (data not shown). Activation of apical caspases independently of effector caspases is a known alternative cell death pathway in *Drosophila* germ cells, but has not yet to our knowledge been described in mammalian germ cells (Yacobi-Sharon *et al.*, 2013).

In this experiment, we also noted occasional spermatids at earlier tubule stages (e.g. stage IV, **Figure 2L**) that appeared to be mis-localised, appearing next to the basement membrane, outside the peripheral ring of meiotic spermatocytes. Sperm heads are generally not seen in this location unless they have been phagocytosed, however these cells were generally negative for both caspase-2 and LAMP2, indicating that they were not yet apoptotic. Since LAMP2 only labels later stages of phagocytosis, we cannot exclude the possibility that these cells were in early stages of phagocytosis, and that phagocytosis in this model may precede the induction of apoptosis.

### Aberrant localisation of Fbxo7^LacZ/LacZ^ condensing spermatids initiates at the onset of cytoplasmic remodelling, from stages I-II onwards

We used periodic acid/Schiff/Hematoxylin (PAS-H) staging to quantify the onset of aberrant localisation of the condensing spermatids in the Fbxo7^LacZ/LacZ^ testes (**Figure 3 A-H**). Here, the PAS staining labels the acrosome, allowing detection of spermatid location and staging. Only very light hematoxylin counterstaining was used to avoid obscuring the PAS signal. For maximal sensitivity in detecting the earliest stages of disorganisation of the seminiferous epithelium, we scored any tubule with even a single spermatid observed at the basement membrane (i.e. appearing to be outside the Sertoli cell tight junctions) as positive. We observed that mis-localisation of late stage spermatids in the Fbxo7^LacZ/LacZ^ testes initiated as early as spermatid step 13-14 (tubule stage I-II), with the proportion of affected tubules apparently peaking at spermatid step 15 (tubule stage IV). From tubule stage VI onwards the phagocytosed spermatid heads were barely visible by PAS-H, most likely due to digestion of the epitopes detected by the PAS stain, and thus the apparent drop-off after stage IV is a technical artefact (note that the dead cells at this stage remain visible via H&E and immunostaining: see **Figures 1**&**2**). This contrasts with the immunostaining data where the mislocalised cells prior to stage VI were LAMP2 and CASP2-negative, but the phagocytosed cells at stages VI-VIII were strongly LAMP2 and CASP2 positive. The two experiments thus probe different aspects of the phenotype: mislocalisation followed by apoptosis. Complete data from the PAS-H cell counting are supplied as **Supplementary Table 2**.

**Figure 3.**
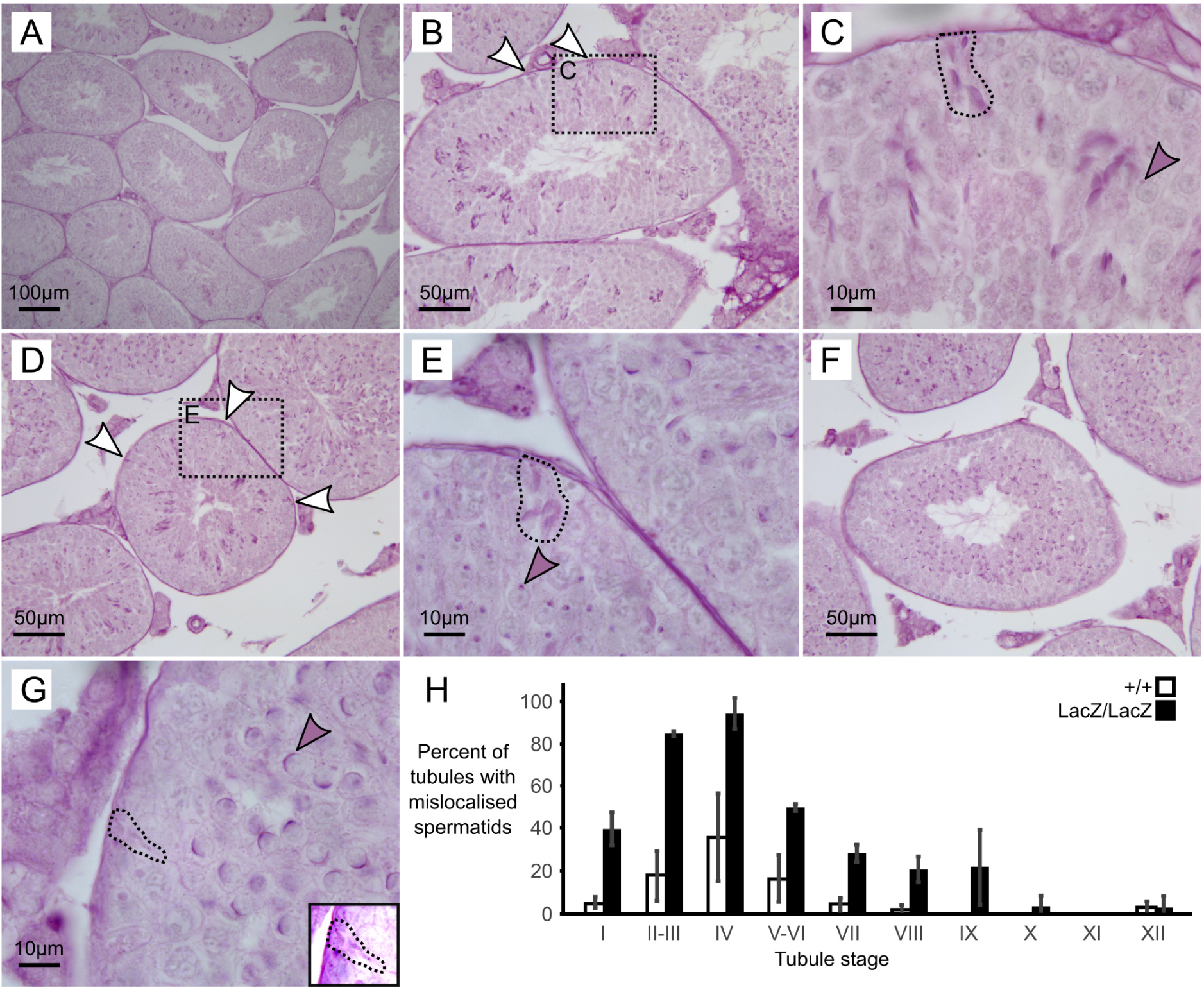
Immunochemical staining of caspase 2 and LAMP2 in Fbxo7^LacZ/LacZ^ testis sections. **A-G:** PAS-H staining of Fbxo7^LacZ/LacZ^ testis sections used to quantitate spermatid mislocalisation. PAS marks the developing acrosome in purple, allowing both identification of the mislocalised elongating spermatids and also tubule staging. **(A)** shows a low magnification view indicating general testis architecture. **(B,D,F)** show complete tubules at stage II, IV and VI respectively, and white arrowheads indicate mislocalised elongating spermatids apposed to the basement membrane of the tubules – these are rarely visible at stage VI. **(C, E, G)** show high magnification images at tubule stages II, IV and VI. Dotted outlines highlight mislocalised elongating spermatids, while shaded arrowheads indicate the developing acrosomes in the round spermatid layer, used to stage the tubules. Mislocalised spermatids were readily detected up to stage IV. After stage IV, the mislocalised cells were still present but began to lose their PAS staining were thus harder to detect. **(G)** inset shows a rare example of a mislocalised cell remaining visible by PAS staining at stage VI. **(H)** Proportion of tubules containing at least one mis-localised cell at each tubule stage in wild type and Fbxo7LacZ/LacZ testes. Error bars indicate standard deviation across replicates (n=3 animals per genotype).

### PI31 expression is reduced in Fbxo7^LacZ/LacZ^ testes, but proteasome activity is unaltered

The basis for sterility in *ntc*-deficient fruit flies is proposed to be the loss of a stabilizing interaction with DmPI31, leading to reduced proteasome activity (Bader *et al.*, 2011). To address whether this relationship is conserved in spermatogenesis in mice, we tested the expression of Fbxo7 and PI31 in lysates made from testes of mature males (**Figure 4A-C**). As expected, the presence of the Fbxo7^LacZ^ allele caused dose-dependent decreases in Fbxo7 expression, as seen by both Western blot and qRT-PCR. We note that for homozygous Fbxo7^LacZ/LacZ^ mice, expression of both mRNA and protein for Fbxo7 was reduced by ~94%. PI31 protein levels were significantly reduced by 30% in Fbxo7^LacZ/LacZ^ testes, while a similar reduction in mRNA levels (23%) was not statistically significant.

**Figure 4.**
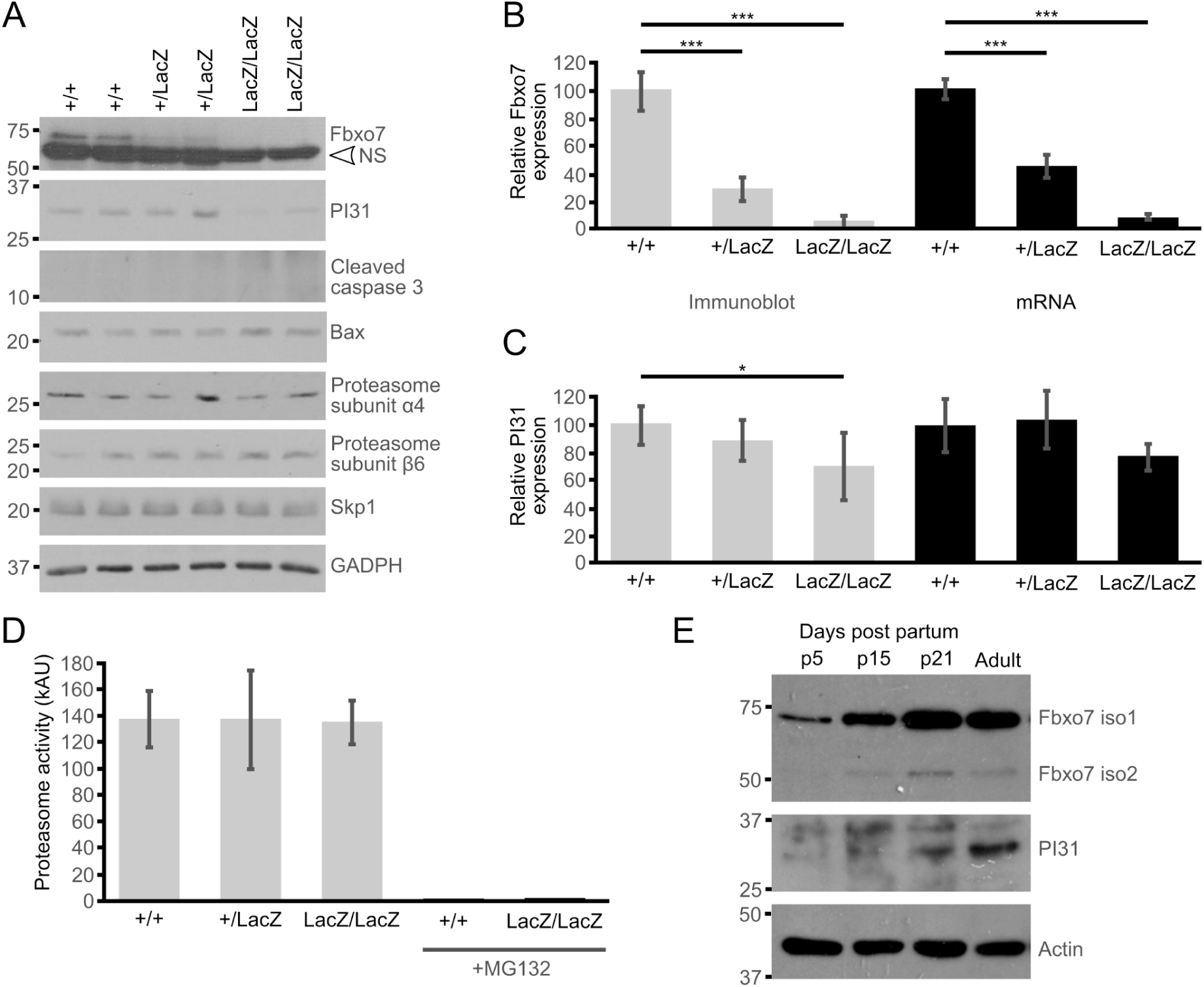
Decreased PI31 levels but normal proteasome activity in Fbxo7^LacZ/LacZ^ testis. **A:** Immunoblot analysis for various proteins in whole testes lysates from WT, heterozygous, and homozygous LacZ mice, as indicated. NS = non-specific band in Fbxo7 Western blot. **B:** Quantification of Fbxo7 expression from immunoblots (relative to GADPH loading control; grey bars) and qPCR relative to three housekeeping genes (cyclophilin, GAPDH, and actin; black bars). Images were analyzed in ImageJ. **C:** Quantification of PI31 expression from immunoblots (relative to GADPH loading control; grey bars) and qPCR of *Psmf1*/PI31 relative to three housekeeping genes (cyclophilin, GAPDH, and actin; black bars). Images were analyzed in ImageJ. **D:** Proteasome activity measured in whole testis extract for each genotype. Treatment with MG132 abolished the signal, confirming the specificity of the assay. **E:** Immunoblot analysis for Fbxo7 and PI31 protein in whole testes lysates from mice harvested at the indicated days postpartum (p).

To test whether the observed reduction in PI31 levels led to decreases in proteasome activity, we conducted proteasome activity assays on whole testes from WT, heterozygous and homozygous Fbxo7^LacZ/LacZ^ males. However, no reduction in proteasome activity was detected (**Figure 4D**). Consistent with these data, we observed no changes in the levels of proteasome subunits among the different WT and mutant testes by Western blot analysis (**Figure 4A**). Attempts to conduct similar proteasome activity assays on elutriated cell populations were inconclusive due to the poor recovery of later stage spermatids from mutant testes (data not shown). These data indicated there were no major alterations in the overall levels of proteasome activity in Fbxo7 deficient testes.

### Spermatoproteasome localisation and histone removal from spermatid chromatin are unaltered in Fbxo7^LacZ/LacZ^ testes

Although spermatoproteasome activity cannot be directly assayed independently of total proteasome activity, the spermatoproteasome has a key role in histone degradation during nuclear elongation (Kniepert and Groettrup, 2014). We therefore considered whether the developmental and spatial distribution of Fbxo7 and/or PI31 are consistent with a role in this process. In developing wild type testes, both Fbxo7 and PI31 were weakly detected at all ages by Western blot, indicating widespread low-level expression in the testis. Both, however, also showed strong upregulation between postnatal day 15 and day 21, concurrent with the first appearance of haploid spermatids in the testis (**Figure 4E**). PI31 was further upregulated between d21 and adult testes, consistent with increased expression in later stage elongating/condensing spermatids. Unfortunately, the available Fbxo7 antibodies do not work for immunohistochemical (IHC) staining in mouse testes. In wild type testes, PI31 was present in the cytoplasm of most cell types, becoming significantly stronger in the cytoplasm of late condensing spermatids from stage V onwards and being retained into the residual bodies shed at stage VIII. In addition to the stage-specific cytoplasmic signal, PI31 also showed nuclear staining specifically in wild type elongating spermatids from stages IX through to XII (**Figure 5**). Thus, there is the potential for Fbxo7 and/or PI31 to regulate the spermatoproteasome during spermatid nuclear remodelling.

**Figure 5.**
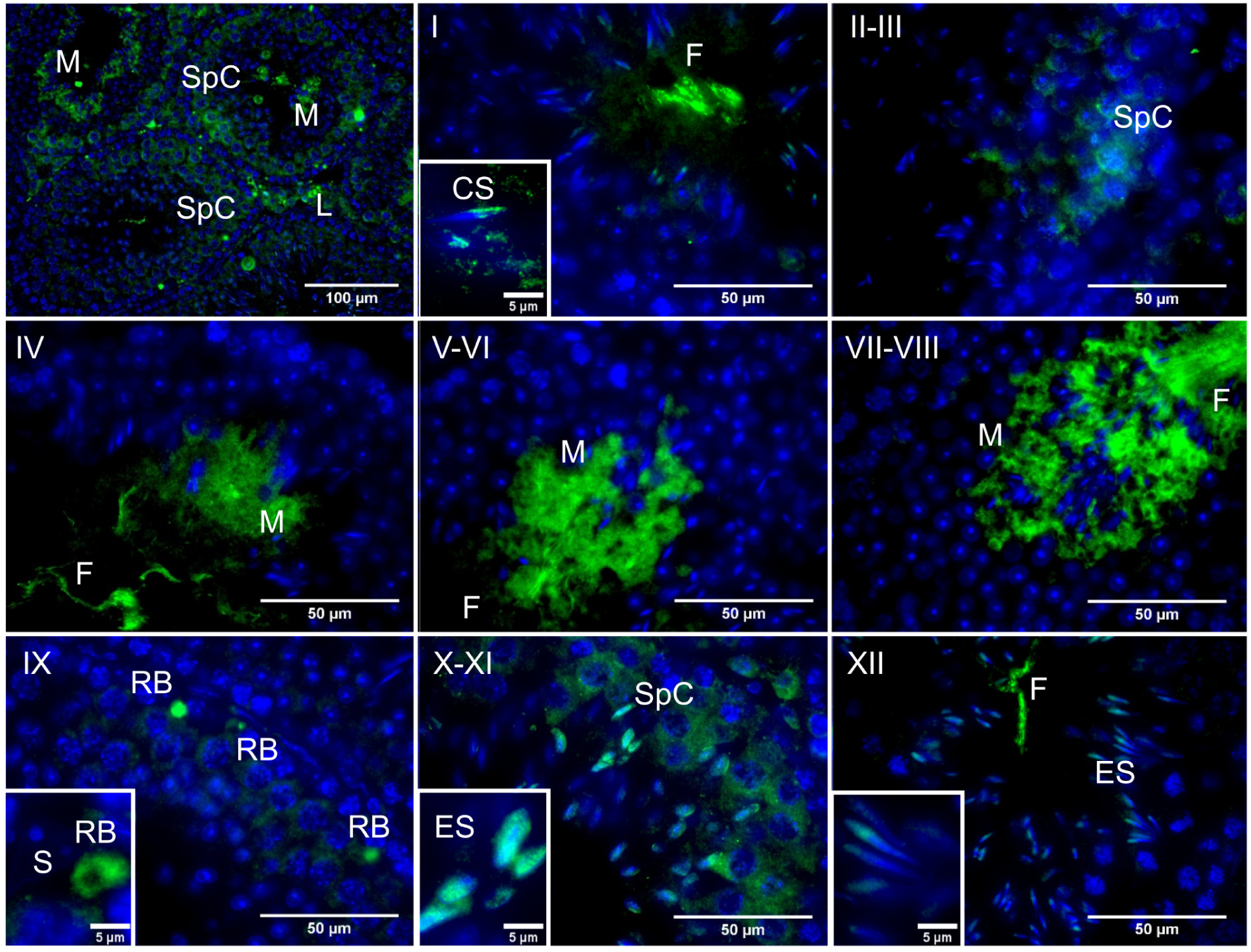
Immunochemical staining of PI31 localisation in wild type testes. Roman numerals indicate tubule stage. PI31 signal is cytoplasmic in Spc / M / RB / L but nuclear in ES and early CS. Key: **SpC** = spermatocyte, **ES** = elongating spermatid, **CS** = early condensing spermatid, **M** = mature or late condensing spermatid, **RB** = residual body, **L** = Leydig cell, **S** = Sertoli cell, **F** = flagellum of mature sperm.

We therefore stained WT and mutant testes for LMP7, a component of the immunoproteasome and spermatoproteasome which is not present in normal proteasomes, to determine whether this was altered by Fbxo7 deficiency (**Figure 6 A**), and for histone H3 to determine whether the dynamics of histone removal was perturbed in the knockout males (**Figure 6 B**). This showed no alteration in spermatoproteasome localisation in the mutant, with nuclear LMP signal being specific to stage IX-XII spermatids in both genotypes. There was also no delay in histone removal in the mutant, with all histone H3 signal being removed from the nucleus by the end of stage XII in both wild type and Fbxo7^LacZ/LacZ^ testes. Taken together with the overall proteasome assay data shown above, we conclude that both spermatoproteasome deficiency in mouse testes. and normal proteasome activity are unaffected by Fbxo7 deficiency in mouse testes.

**Figure 6.**
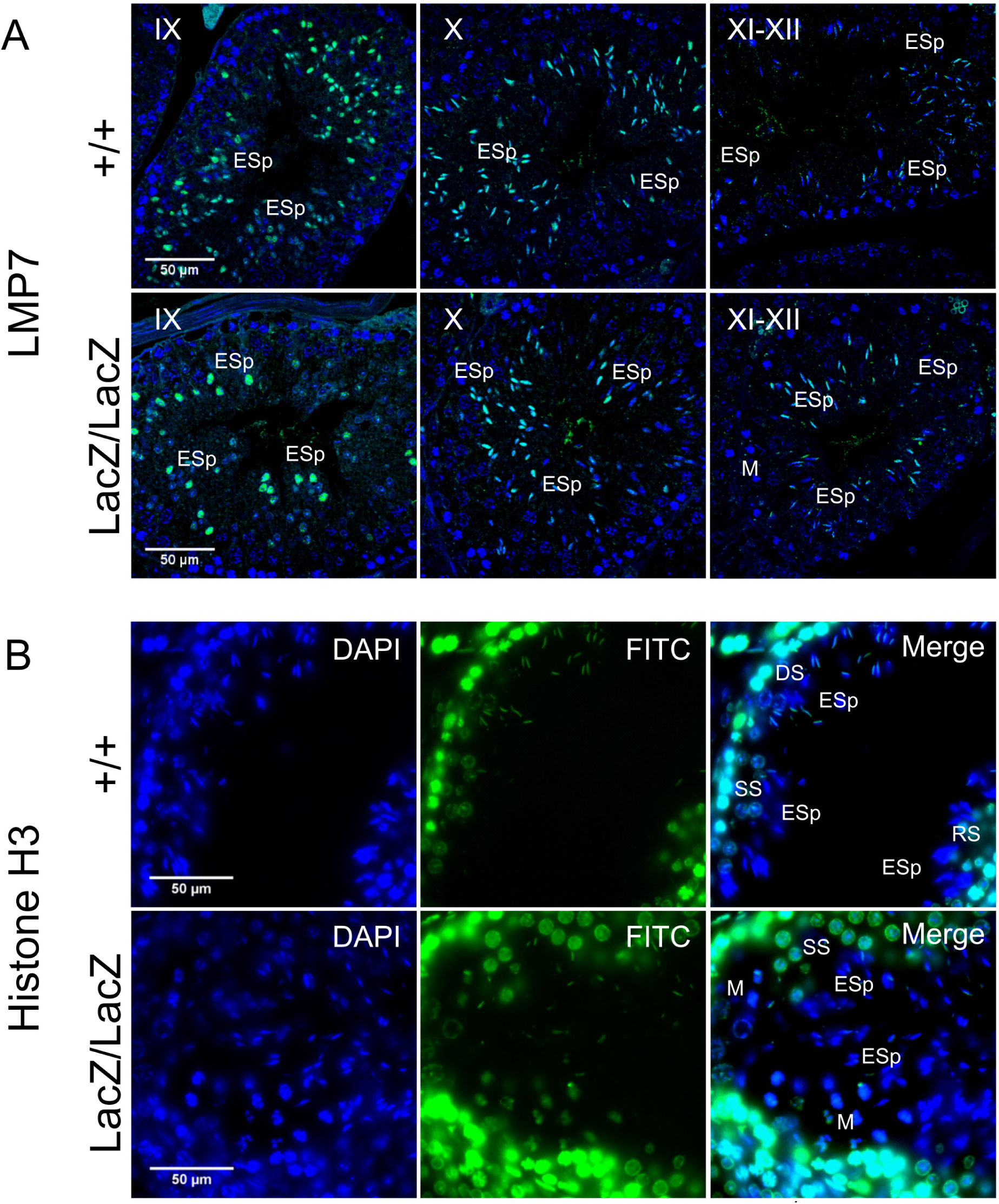
Immunochemical staining of LMP7 and histone H3 in Fbxo7^LacZ/LacZ^ testis sections. **A:** Immunochemical staining of LMP7 (FITC, green) in wild type and Fbxo7^LacZ/LacZ^ testes with DAPI (blue) nuclear counterstain. Roman numerals indicate tubule stage. In both genotypes, nuclear LMP7 expression is first seen in early ES at mid-stage IX as the nuclei begin to elongate. This nuclear expression is highest at stage X, and then is lost during stage XI-XII as nuclei complete elongation. **B:** Immunochemical staining of histone H3 (FITC, green) in wild type and Fbxo7^LacZ/LacZ^ testes with DAPI (blue) nuclear counterstain. The wild type tubule shown is in transition between stages, with early stage XII (DS next to ESp) at upper left, mid stage XII (SS next to ESp) at lower left and stage XII/I border (M and RS next to ESp) at lower right. Nuclear H3 signal in ESp is still present at early stage XII, is restricted to the posterior of the nucleus in mid stage XII, and entirely lost by stage I. The Fbxo7^LacZ/LacZ^ tubule shown is in mid stage XII, and the H3 signal in the ESp is absent or restricted to the posterior extremity of the nucleus, confirming the kinetics and spatial pattern of H3 removal are indistinguishable between genotypes. Key: **DS** = diplotene spermatocytes, **SS** = secondary spermatocytes, **M** = metaphase figures, **RSp** = round spermatids, **ESp** = elongating spermatids.

## Discussion

### The mammalian phenotype associated with Fbxo7 deficiency

Spermiogenesis is a multi-step process that transforms morphologically simple round spermatids into highly-specialised mature sperm. It occurs in four successive phases, namely; **a)** nuclear elongation and replacement of histones by transition proteins in tubule stages IX to XII, spermatid step 9-12), **b)** protamination of sperm chromatin and cytoplasmic reduction by ~50% in tubule stages I to VI, spermatid steps 13-15, **c)** migration of the mature spermatids to the tubule lumen in tubule stage VII, spermatid early step 16, **d)** spermiation, the release of fully-formed sperm at tubule stage VIII, spermatid late step 16. In Fbxo7^LacZ/LacZ^ males, we observe a peculiar and very specific phenotype consisting of mass phagocytosis of condensing spermatids, occurring after nuclear elongation and prior to migration of spermatids back to the tubule lumen. This coincides with the cytoplasmic remodelling of the spermatids during steps 13-15, equivalent to the individuation stage of *Drosophila* sperm development. The requirement for Fbxo7 at this stage thus appears to be strictly conserved from fruit flies to mammals.

The nature of the sterility phenotype in Fbxo7^LacZ/LacZ^ males is to our knowledge unprecedented in a mammalian system. Various null mutants with defects in cytoplasmic remodelling have previously been described, including Sept4, Spem1, Capza3, and Ube2j1 mutants. These all show either no phagocytosis or only limited phagocytosis of elongating spermatids during the first half of the cycle, followed by spermiation failure and spermatid retention at the lumen into stage IX and beyond (Kissel et al., 2005; Zheng et al., 2007; Geyer et al., 2009; Koenig et al., 2014). Further mutants have been described that shows a step 13 block to spermatid development without defects in cytoplasmic remodelling, including the Bclw and Brd7 null mutants. In these mice, the arrested step 13 spermatids degenerate while still near the tubule lumen, followed by phagocytosis of large symplasts and other cell debris (Russell et al., 2001; Wang et al., 2016). In stark contrast to both of the above types of mutant, the Fbxo7^LacZ/LacZ^ males showed complete phagocytosis of all developing spermatids with no detectable symplast formation, sloughing of degenerating cells into the lumen, or spermiation failure.

### How are the germ cells eliminated in Fbxo7^LacZ/LacZ^ testes?

In Fbxo7^LacZ/LacZ^ testes, mis-localised condensing spermatids are visible by PAS-H staining at the basement membrane from tubule stage ~I-II onwards, and by stage IV almost 100% of tubules have mis-localised cells. At these early stages, however, mis-localised spermatids are negative for caspase-2 and LAMP-2, and retain their acrosomes (i.e. they are stainable by PAS), suggesting that they are not yet apoptotic and/or that phagocytic degradation has not yet initiated. By stage VI, however, the cells have lost their normal orientation, become positive for caspase 2 and LAMP-2, and karyolysis has initiated.

One possible scenario is that the early stages of mis-localisation represent an abnormal deepening of the Sertoli cell crypts in mutant testes, and that the condensing spermatids have not yet been phagocytosed at this point. The trafficking of spermatids into and out of Sertoli cell crypts is governed by dynein-coupled motion of a specialised adherens junction complex between germ cell and Sertoli cell, known as the <u>a</u>pical <u>e</u>ctoplasmic <u>s</u>pecialisation (AES). If the AES is dissolved prematurely, the Sertoli cell may be unable to eject the spermatids from the crypts, leading ultimately to their death and phagocytosis.

An alternative scenario is that the phagocytosis initiates at stage I-II due to defects in germ cell remodelling, but that the engulfed cells remain alive and non-apoptotic for a short period after being phagocytosed. That is, the developing spermatids are “eaten alive” some time before finally dying. In this case, this would represent death by primary phagocytosis or “phagoptosis”, a relatively newly-identified manner of cell elimination (Brown and Neher, 2012). Phagoptosis of elongating spermatids in Fbxo7^LacZ/LacZ^ testes would explain the absence of symplasts and the lack of sloughing of dead cells into the lumen, since in the case of phagoptosis, the eliminated cells would remain alive until they have already become trapped inside phagocytic vacuoles.

A formal test of these hypotheses will require electron microscopic investigation to resolve the relevant intracellular machinery and distinguish between deep invagination of elongating spermatids into Sertoli cell crypts versus early stages of engulfment during phagocytosis.

### Are the consequences of Fbxo7 deficiency mediated by PI31 / proteasome regulation?

The phenotype of *ntc* mutant testes in *Drosophila* has been ascribed to an interaction with PI31 that regulates proteasome activity in spermatids; however, this is not conclusively proven as transgenic restoration of PI31 levels in *ntc* testes was unable to correct the defect in spermatid individualisation (Bader *et al.*, 2011). We and others have previously shown that the stabilising interaction between Fbxo7 and PI31 is conserved. Consistent with this, we show here that PI31 levels are reduced in Fbxo7^LacZ/LacZ^ testes. In wild type testes, we show that PI31 and LMP7 are both present in the nucleus of step 10-12 spermatids, indicating that spermatoproteasomes most likely play a role in nuclear elongation. PI31 then shifts to the cytoplasm of step 13-16 spermatids at tubule stages I-VIII.

The loss of germ cells in the knockout is thus coincident with this shift in PI31 localisation from the nucleus to the cytoplasm. It is possible therefore that PI31 expression in the cytoplasm is necessary during spermatid remodelling, and Fbxo7 is required to stabilise PI31 in the cytoplasm. Set against this, however, is the fact that cytoplasmic PI31 is highest in tubule stages VI-VIII, by which time the Fbxo7^LacZ/LacZ^ spermatids are already dead and digested, and very low in stages II-IV when spermatid mis-localisation initiates in Fbxo7^LacZ/LacZ^ testes. Moreover, both the mRNA and protein product for PI31 declined to a similar degree, suggesting that the decrease in both may be a secondary consequence of the loss of late stage condensing spermatids in mutant testes rather than a change in the stability of PI31. Overall, therefore, the data on PI31 expression and localisation suggests that in mouse, the consequences of Fbox7 deficiency for male fertility are not mediated by a secondary deficiency for PI31.

Although we could not directly measure spermatoproteasome activity, whole-testis proteasome activity showed no changes in Fbxo7^LacZ/LacZ^ testes. Other proteasome-related knockout males (PA200, PA28γ) have defects in multiple stages of germ cell development, both pre- and post-meiotic. The post-meiotic phenotypes of these knockout models include delayed histone replacement during nuclear remodelling and delayed spermiation, neither of which was seen in Fbxo7^LacZ/LacZ^ testes (Qian et al., 2013; Huang et al., 2016). Moreover, unlike Fbxo7^LacZ/LacZ^ males, even the PA200/PA28γ double knockout males were able to produce substantial numbers of morphologically normal sperm in their epididymis (Huang *et al.*, 2016). A knockout of the spermatoproteasome-specific subunit PSMA8 has recently been shown to lead to meiotic abnormalities and early spermatid arrest [Gómez Hernández et al, unpublished data, preprint available at https://www.biorxiv.org/content/early/2018/08/03/384354], unlike the late stage spermatid loss described in the present study. The Fbxo7^LacZ/LacZ^ male sterility phenotype thus differs from all other existing mouse knockouts related to proteasome function at both the histological and molecular levels, and this argues against proteasome insufficiency being causative for its phenotypes.

### Potential non-proteasomal pathways regulated by Fbxo7 that may lead to male sterility

Fbxo7 is a multi-functional protein which impacts upon many pathways, such as cytoplasmic remodelling, mitophagy and cell cycle regulation, which are key process during the production of specialized cell types. Fbxo7 is required for Parkin-mediated mitophagy, a process that requires the fragmentation and engulfment of depolarized regions of the mitochondrial network (Burchell *et al.*, 2013), and interestingly, both the *nutcracker* and *parkin* null flies show male sterility, with a specific defect during sperm individuation (Greene et al., 2003; Bader et al., 2010). In the *parkin* null, a specialized mitochondrial aggregate present in insect sperm, known as the Nebenkern, failed to form, suggesting the rearrangement of mitochondria was part of the underlying mechanism of sterility. However, the mouse parkin null mutant is fertile with no known effects on germ cell development (Itier *et al.*, 2003), and thus the sterility of Fbxo7 mutant males is unlikely to relate directly to its interaction with Parkin.

One possible explanation is that Fbxo7 is not only involved in mitophagy through the PINK1/Parkin pathway, but also more generally with specialized cytoplasmic remodelling. In a similar vein to the sperm maturation defects we report here, Fbxo7 has also been shown to be required during the final maturation steps of erythrocytes, and we have previously reported the Fbxo7^LacZ/LacZ^ mice are anaemic due to delayed mitophagy and defects in exiting cell cycle (Randle and Laman, 2016). Importantly, during this maturation step, macrophages in erythroblast islands phagocytose the shed organelles from maturing reticulocytes (Ovchynnikova et al., 2018; Zhang et al., 2015; Geminard et al., 2002), a process requiring the coupling of the autophagy and exocytosis pathways (Mankelow *et al.*, 2015). Could this coupling be coordinated by Fbxo7? If so, then in a testis context, Fbxo7 may enable the fragmentation and isolation of portions of the spermatid cytoplasm to allow phagocytosis by Sertoli cells. In the absence of Fbxo7, failure to correctly package spermatid cytoplasm for elimination could instead lead to wrongful engulfment of complete cells and phagoptotic cell death.

As a third alternative but non-exclusive possibility, we note that the dead cells at tubule stage VI were strongly positive for caspase 2. TRAF2, a target of Fbxo7 ubiquitination (Kuiken *et al.*, 2012), has recently been shown to bind to active caspase 2 dimers and ubiquitinate it to stabilize the activated complex (Robeson *et al.*, 2018). Consequently, Fbxo7 deficiency could lead to over-activity of TRAF2 and subsequently signal the activation of caspase 2 precipitating germ cell death. Identifying the substrates of Fbxo7 underlying the unique phenotypes reported here is an area of future research.

### Conclusion

*Fbxo7* deficient mice exhibit a novel sterility phenotype unlike any previously described, in that total death and phagocytosis of all condensing spermatids occurs in the absence of typical hallmarks of spermatid apoptosis such as symplast formation and cell sloughing. The mislocalisation of elongating spermatids initiates substantially before the appearance of markers of apoptosis and phagocytosis, indicating that aspects of spermatid trafficking into and out of Sertoli cell crypts may also be perturbed in these males. These males thus provide a new model of late spermiogenic failure, and an exciting new avenue to investigate cell remodelling, tissue remodelling and apoptosis in germ cell development.

## Materials and Methods

### Mice

Mice used in this study are *Fbxo7*^*LacZ*^ mice (*Fbxo7*^*tm1a(EUCOMM)Hmgu*^ on a C57BL/6J background) and experiments involving them were performed in accordance with the UK Animals (Scientific Procedures) Act 1986 and ARRIVE guidelines. Mice were housed in individually ventilated cages with unrestricted access to food and water, and 12-hour day-night cycle. Animal licences were approved by the Home Office and the University of Cambridge’s Animal Welfare & Ethical Review Body Standing Committee. Experiments were performed under the Home Office licences PPL 80/2474 and 70/9001 (HL).

### Tissue processing and immunohistochemistry

Testes were fixed in Bouin’s fixative at 4^0^C overnight and embedded in paraffin. Sections were subjected to standard methods of hematoxylin/eosin or periodic acid/Schiff/hematoxylin staining for histological examination. For immunohistochemical studies, sections were de-paraffinised in xylene and rehydrated through a graded ethanol series prior to blocking and antibody staining steps. Details of primary and secondary antibody concentrations are given in **Supplementary Table 1**. Antibody-stained sections were counterstained with DAPI and visualised via epifluorescence (PI31, Histone H3) or confocal fluorescence microscopy (Caspase 2, Lamp-2, LMP7). In some experiments, fluorescently conjugated peanut agglutinin from Arachis hypogaea (PNA-lectin) was included during the secondary antibody incubation step to visualise acrosomal morphology and facilitate staging of seminiferous tubules.

### Sperm morphometric analysis

Sperm were collected from two Fbxo7^LacZ/LacZ^, three Fbxo7^LacZ/+^, and two wild type males. The vasa deferentia and caudae epididymides were dissected from each animal, and the contents extracted into 1ml PBS. Sperm were transferred to a microfuge tube, and tissue clumps were allowed to settle. Then sperm were transferred to a new tube and pelleted at 500g for 5mins. The supernatant was removed, and the sperm fixed dropwise with 3:1 methanol-acetic acid. Sperm were again pelleted at 500g for 5mins, and washed in fixative twice more. Samples were stored at −20°C. Fixed sperm nuclei were stained with DAPI and imaged using a 100x objective on a Nikon Microphot SA epifluorescence microscope equipped with a cooled CCD camera and appropriate filters. Images were captured using SmartCapture 2, exported in 8-bit tiff format and analysed using the automated morphometric software Nuclear Morphology Analysis v1.13.7 (https://bitbucket.org/bmskinner/nuclear_morphology/wiki/Home) (Skinner et al., 2019). Hierarchical clustering was performed on nuclear shapes to group them into morphological categories, and the proportion of cells from each genotype in each category was calculated. The total numbers of nuclei analysed for each genotype were 453 for Fbxo7^LacZ/LacZ^, 1225 for Fbxo7^LacZ/+^ and 756 for wild type.

### Flow cytometry

Single-cell suspensions from whole testis tissue were pelleted by centrifugation, and resuspended in 1mL of ice cold 80% ethanol/PBS while vortexing to disperse clumps. The suspended cells were fixed for at least 1 hour at 4°C. After fixation, cells were collected by centrifugation and washed once in PBS. The washed cell pellet was resuspended in 1mL of a solution of 50μg/mL propidium iodide (PI) staining and 10μg/mL DNase-free RNase, and incubated for 10 minutes at 37°C, prior to analysis by flow cytometry (Beckman-Coulter, Inc.).

### Lysis and immunoblotting

Whole testis tissue were lysed in RIPA buffer (50mM Tris-HCl pH 7.6, 150mM NaCl, 1% NP-40, 0.1% SDS, 0.1% Na deoxycholate, 1x protease inhibitors, 1mM PMSF, 10mM sodium fluoride, 1mM sodium orthovanadate) (all from Sigma), and incubated on ice for 30 min with occasional vortexing. Cell debris was pelleted by centrifugation at 16,000g for 10 min at 4°C, and the supernatant retained. Protein concentration was determined via 96-well BCA assay (Pierce). Sample concentrations were standardised by dilution with lysis buffer. For western blot, samples were mixed with equal volumes of 2x Laemmli buffer, denatured (95°C, 5mins), separated via SDS polyacrylamide gel electrophoresis (SDS-PAGE), and transferred onto polyvinylidenefluoride (PVDF) membrane (Millipore) using a semi-dry transfer system (Biorad). Membranes were blocked for one hour with 5% non-fat, milk powder/PBS-Tween 20 (0.05%) (PBS-T), and then probed with primary antibody overnight at 4°C in 5% non-fat, milk powder/PBS-T. Membranes were washed in PBS-T and incubated with the appropriate HRP-conjugated secondary antibody in 5% non-fat, milk powder/PBS-T followed by further washes, and detection of HRP bound protein using enhanced chemiluminescence (ECL, GE Healthcare) and exposure onto X-ray film (Konica Minolta). Signal was quantified and normalised using ImageJ software (NIH, Maryland).

### mRNA isolation and qRT-PCR

Tissue was homogenised in 350 µl RLT buffer with β-mercaptoethanol and RNA isolated using RNeasy Plus kit (Qiagen) as per the manufacturer’s recommendations. One microgram of mRNA was converted to cDNA using Quantitect reverse transcriptase (Qiagen), and then diluted 1:10 for subsequent qRT-PCR analysis using SYBR Green JumpStart Taq (Sigma) on a CFX Connect Real-Time PCR machine (Biorad). The following primers for *Fbxo7* (5’-CGCAGCCAAAGTGTACAAAG; 3’-AGGTTCAGTACTTGCCGTGTG) and *Psmf1*(PI31) (5’-CAATCATGCCACCTCTCTGA; 3’-CCGTCCTCATACTAGCAGGC) were used. qRT-PCR reactions were as follows: 95°C for 5 min then 45 cycles of 95°C for 30 s, 60°C for 30 s, 72°C for 30 s, followed by melt curve analysis to confirm a single PCR product was made. Relative gene expression was determined using relative standard curve method, data was normalised to three housekeeping genes (*Ppai, Gapdh, Actb*) as previously described [see (Meziane et al., 2011; Birkenfeld et al., 2011) for primer sequences], and expressed relative to WT levels.

### Counting of mis-localised cells at different tubule stages

Tubules were staged using periodic acid/Schiff staining to visualise the stages of acrosomal development (Russell et al., 93 A.D.; OAKBERG, 1956). Every tubule in a complete testis cross-section was staged for three replicate males per genotype, by an observer blinded to the sample identity. Tubules were scored as positive if there were any mis-located elongating spermatid heads detected beyond the Sertoli cell tight junctions, within the outermost layer of nuclei in the tubule, and negative if there were no elongating spermatid heads within this layer. Tubules were also scored for the presence of “graveyards”, defined as 2 or more mis-localised elongated spermatids enveloped by a single Sertoli cell. These definitions were chosen to maintain consistency across the seminiferous cycle, and their sensitivity is discussed in the main text.

### Measurement of proteasome activity

*Assays* for proteasome activity were performed using the Proteasome-Glo™ Chymotrypsin-Like Cell-Based Assay Kit (Promega) according to the manufacturer’s protocol. Briefly, testes from 10 week old mice were harvested and lysed in buffer (20 mM Hepes pH 7.6, 150 mM NaCl, 10% glycerol, 1% Triton X-100, 2 mM EDTA, 1 mM DTT) using a Dounce homogeniser in 10X the volume/weight of tissue. Protein concentration was measured, and lysates were diluted so that 100μg of protein in 100μl volume/well was loaded into a 96-well plate, and samples were plated in triplicate. Where applicable, samples were pre-incubated with MG-132 for 30 minutes prior to the addition of reagents, with protease inhibitors (Na_2_VO_4_, NaF, PSMF), or without inhibitors. Samples were equilibrated to RT for 15 minutes and then 100μl of assay reagent was added. After a 10 minute incubation, luminescence was measured in triplicate.

## Acknowledgements

This work was supported by the BBSRC (BB/J007846/1 to HL, BB/N000463/1 to PE, BB/N000129/1 to NA).

## Declarations of Interests

No competing interests declared.

## Supplementary Data

**Supplementary Figure 1.**
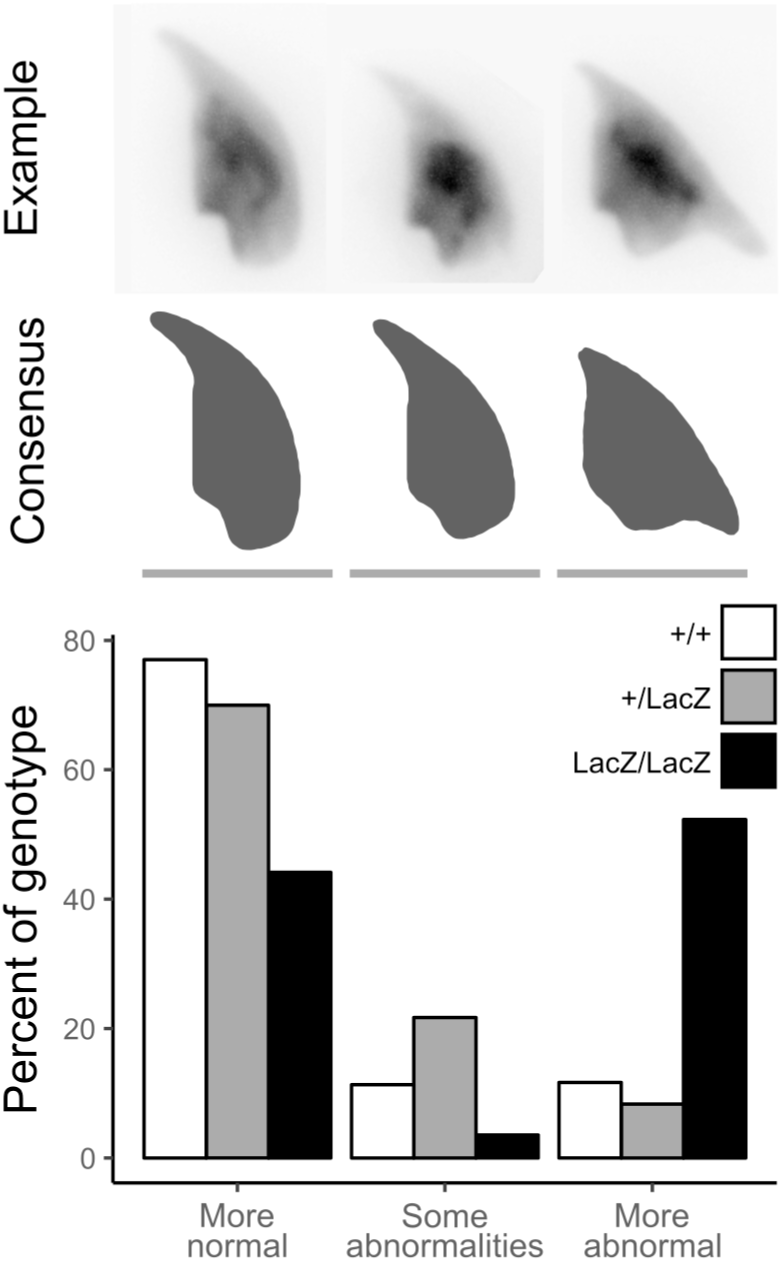
Sperm shape abnormalities are overrepresented in Fbxo7^LacZ/LacZ^ animals compared to wild type and heterozygous animals. Sperm from all genotypes were clustered according to shape into three categories of normal, somewhat abnormal and severely deformed. Upper panel; representative DAPI-stained sperm nuclei from each cluster; middle panel; consensus shape of the cluster; lower panel; percent of each genotype within the cluster. The majority of the Fbxo7^LacZ/LacZ^ sperm have severe abnormalities, compared with less than 10% of wild type or heterozygote sperm.

**Supplementary Figure 2.**
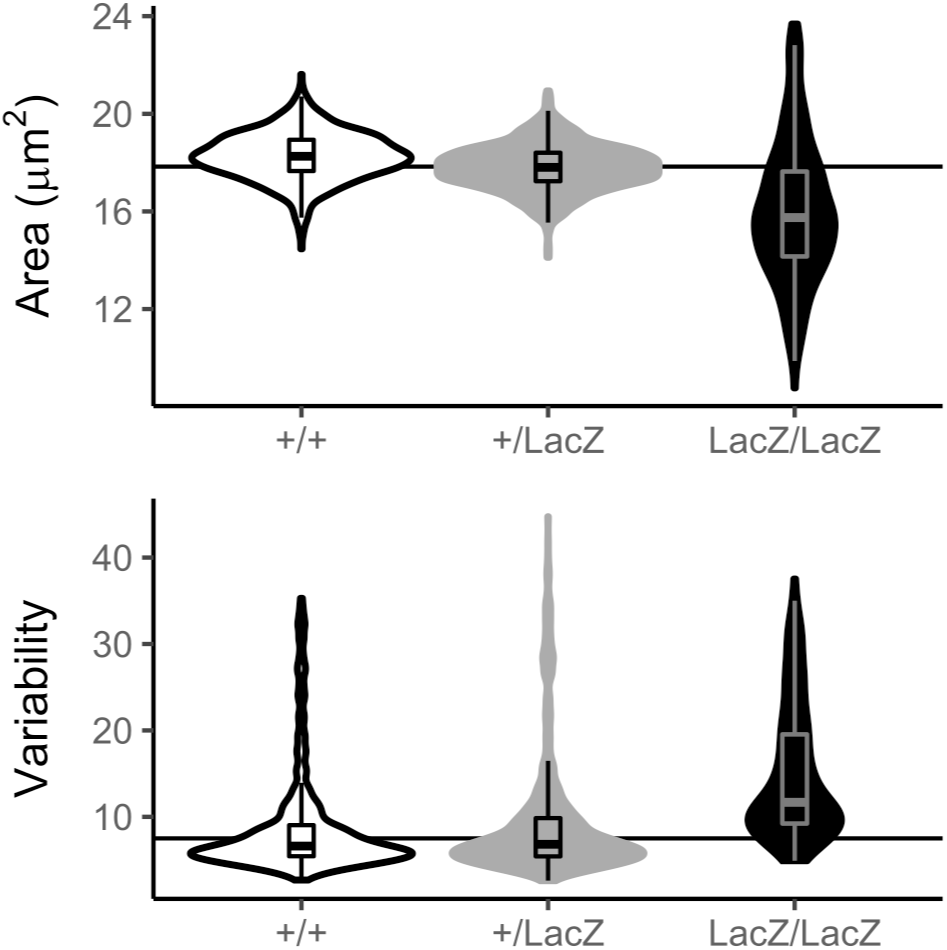
Sperm from Fbxo7^LacZ/LacZ^ animals are mostly smaller than sperm from wild type and heterozygous animals, with higher variability (measured as the difference between each nucleus’ shape and the median shape).

**Supplementary Table 1:**
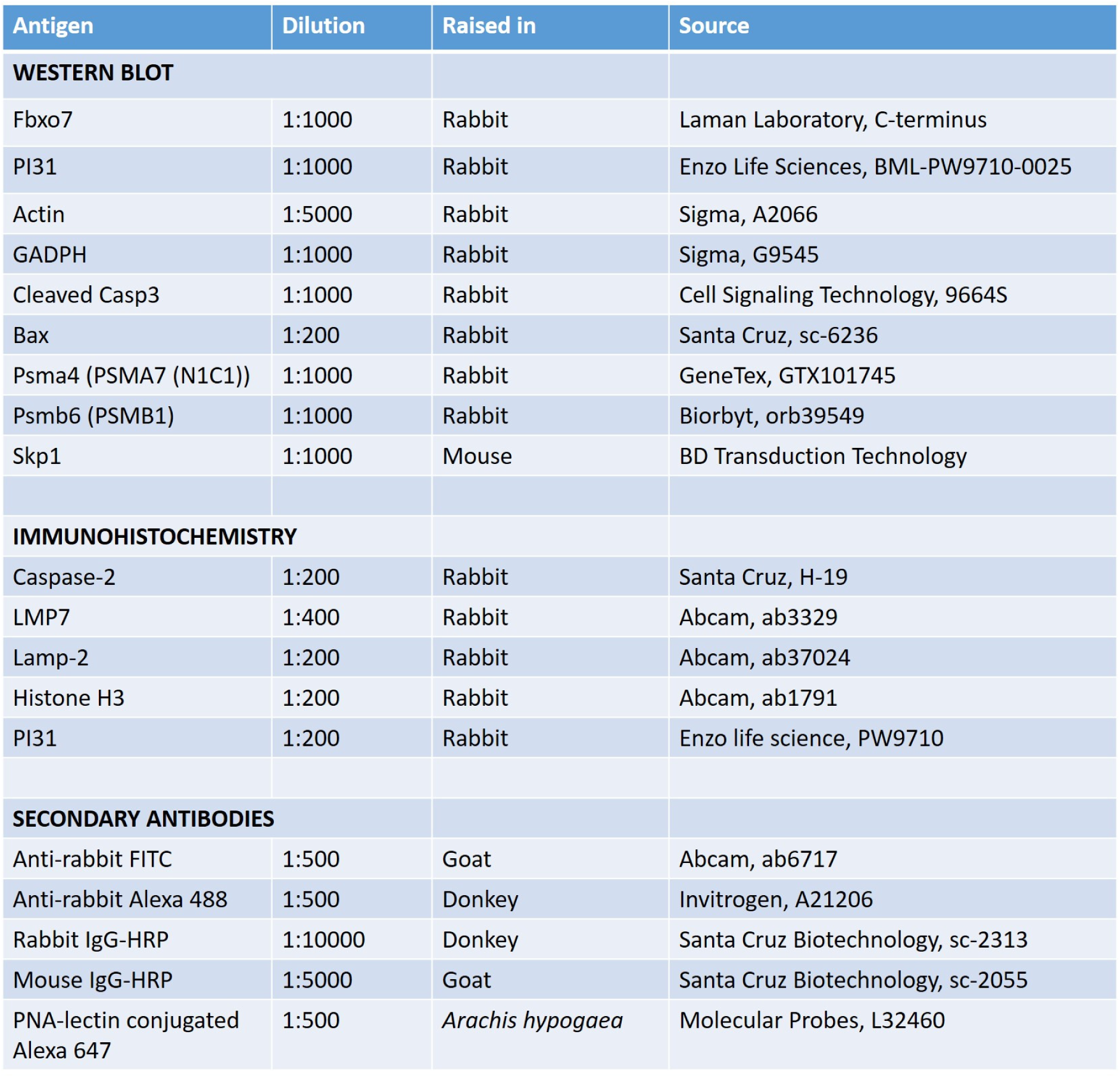
Antibodies used in this study

