## Supplementary Table 2 for "A conserved requirement for *Fbxo7* during male germ cell cytoplasmic remodelling"

**All tubules were examined for a complete testis cross section for three males from each genotype**
**sgl = tubules with one or more singleton spermatid heads located at the basement membrane**
**gy = tubules with one or more "graveyards" consisting of 2 or more mislocalised spermatids lying within the same Sertoli cell.**

### Aggregate figures

**RAW NUMBERS**

| | WT | | | | LacZ/LacZ | | | | |
|---|---|---|---|---|---|---|---|---|---|
| Stage | Tubules with gy | Tubules with sgl | Tubules with any | Total tubules | Tubules with gy | Total gy | Tubules with sgl | Tubules with any | Total tubules |
| I | 1 | 3 | 4 | 85 | 10 | 19 | 12 | 22 | 55 |
| II-III | 7 | 4 | 11 | 61 | 34 | 45 | 17 | 51 | 60 |
| IV | 14 | 5 | 19 | 53 | 18 | 34 | 16 | 34 | 36 |
| V-VI | 7 | 4 | 11 | 67 | 23 | 37 | 23 | 46 | 92 |
| VII | 0 | 2 | 2 | 48 | 10 | 25 | 8 | 18 | 63 |
| VIII | 0 | 1 | 1 | 52 | 7 | 15 | 2 | 9 | 43 |
| IX | 0 | 0 | 0 | 56 | 3 | 4 | 9 | 12 | 55 |
| X | 0 | 0 | 0 | 36 | 1 | 1 | 0 | 1 | 32 |
| XI | 0 | 0 | 0 | 40 | 0 | 0 | 0 | 0 | 29 |
| XII | 0 | 1 | 1 | 36 | 1 | 1 | 0 | 1 | 35 |
| | | | | 534 | | | | | 500 |

**NORMALISED PERCENT PER TUBULE STAGE**

| | WT | | | | LacZ/LacZ | | | | |
|---|---|---|---|---|---|---|---|---|---|
| Stage | % Tubules with gy | % Tubules with sgl | % Tubules with any | Total % Tubules ( | % Tubules with gy ( | gy per tul | % Tubules with sgl ( | % Tubules with any (LacZ/LacZ) | Total % Tubules (LacZ/LacZ) |
| I | 0.01 | 0.04 | 0.05 | | 0.18 | 0.35 | 0.22 | 0.40 | |
| II-III | 0.11 | 0.07 | 0.18 | | 0.57 | 0.75 | 0.28 | 0.85 | |
| IV | 0.26 | 0.09 | 0.36 | | 0.50 | 0.94 | 0.44 | 0.94 | |
| V-VI | 0.10 | 0.06 | 0.16 | | 0.25 | 0.40 | 0.25 | 0.50 | |
| VII | 0.00 | 0.04 | 0.04 | | 0.16 | 0.40 | 0.13 | 0.29 | |
| VIII | 0.00 | 0.02 | 0.02 | | 0.16 | 0.35 | 0.05 | 0.21 | |
| IX | 0.00 | 0.00 | 0.00 | | 0.05 | 0.07 | 0.16 | 0.22 | |
| X | 0.00 | 0.00 | 0.00 | | 0.03 | 0.03 | 0.00 | 0.03 | |
| XI | 0.00 | 0.00 | 0.00 | | 0.00 | 0.00 | 0.00 | 0.00 | |
| XII | 0.00 | 0.03 | 0.03 | | 0.03 | 0.03 | 0.00 | 0.03 | |

**STANDARD DEVIATION OF PERCENTAGES**

| | StDev (WT) | | | | StDev (LacZ/LacZ) | | | | |
|---|---|---|---|---|---|---|---|---|---|
| Stage | % Tubules with gy | % Tubules with sgl | % Tubules with any | Total % Tubules ( | % Tubules with gy ( | gy per tul | % Tubules with sgl ( | % Tubules with any (LacZ/LacZ) | Total % Tubules (LacZ/LacZ) |
| I | 0.023094011 | 0.007583554 | 0.027047816 | | 0.07889238 | 0.18527 | 0.087128453 | 0.0773294 | |
| II-III | 0.123260663 | 0.016159163 | 0.119947331 | | 0.036899053 | 0.18779 | 0.043822521 | 0.012262313 | |
| IV | 0.16938438 | 0.041342453 | 0.208725554 | | 0.095470327 | 0.24855 | 0.157288217 | 0.072168784 | |
| V-VI | 0.070299455 | 0.052486388 | 0.115439382 | | 0.15997308 | 0.30684 | 0.146316646 | 0.017337503 | |
| VII | 0 | 0.038570443 | 0.038570443 | | 0.097837218 | 0.36752 | 0.074435721 | 0.040357637 | |
| VIII | 0 | 0.02220578 | 0.02220578 | | 0.031264424 | 0.29479 | 0.033599898 | 0.063898256 | |
| IX | 0 | 0 | 0 | | 0.047838998 | 0.06009 | 0.142046495 | 0.180282805 | |
| X | 0 | 0 | 0 | | 0.052486388 | 0.05249 | 0 | 0.052486388 | |
| XI | 0 | 0 | 0 | | 0 | 0 | 0 | 0 | |

| XII | 0 | 0.036084392 | 0.036084392 | | 0.057735027 | 0.05774 | 0 | 0.057735027 |
|---|---|---|---|---|---|---|---|---|

### Individual animal counts

| | WT (12/122/951) | | | | LacZ/LacZ (912/122/950) | | | | |
|---|---|---|---|---|---|---|---|---|---|
| | Tubules with gy | Tubules with sgl | Tubules with any | Total tubules | Tubules with gy | Total gy | Tubules with sgl | Tubules with any | Total tubules |
| I | 1 | 1 | 2 | 25 | 3 | 7 | 3 | 6 | 12 |
| II-III | 0 | 2 | 2 | 27 | 8 | 13 | 3 | 11 | 13 |
| IV | 2 | 1 | 3 | 20 | 9 | 17 | 5 | 14 | 16 |
| V-VI | 4 | 2 | 6 | 22 | 12 | 21 | 2 | 14 | 27 |
| VII | 0 | 1 | 1 | 23 | 6 | 11 | 1 | 7 | 23 |
| VIII | 0 | 1 | 1 | 26 | 3 | 6 | 1 | 4 | 19 |
| IX | 0 | 0 | 0 | 21 | 2 | 2 | 4 | 6 | 21 |
| X | 0 | 0 | 0 | 17 | 1 | 1 | 0 | 1 | 11 |
| XI | 0 | 0 | 0 | 20 | 0 | 0 | 0 | 0 | 10 |
| XII | 0 | 1 | 1 | 16 | 1 | 1 | 0 | 1 | 10 |
| | | | | 217 | | | | | 162 |
| | % Tubules with gy | % Tubules with sgl | % Tubules with any | Total % Tubules ( | % Tubules with gy (l | gy per tul | % Tubules with sgl (l | % Tubules with any (LacZ/LacZ) | Total % Tubules (LacZ/LacZ) |
| I | 0.04 | 0.04 | 0.08 | | 0.25 | 0.58333 | 0.25 | 0.5 | |
| II-III | 0 | 0.074074074 | 0.074074074 | | 0.615384615 | 1 | 0.230769231 | 0.846153846 | |
| IV | 0.1 | 0.05 | 0.15 | | 0.5625 | 1.0625 | 0.3125 | 0.875 | |
| V-VI | 0.181818182 | 0.090909091 | 0.272727273 | | 0.444444444 | 0.77778 | 0.074074074 | 0.518518519 | |
| VII | 0 | 0.043478261 | 0.043478261 | | 0.260869565 | 0.47826 | 0.043478261 | 0.304347826 | |
| VIII | 0 | 0.038461538 | 0.038461538 | | 0.157894737 | 0.31579 | 0.052631579 | 0.210526316 | |
| IX | 0 | 0 | 0 | | 0.095238095 | 0.09524 | 0.19047619 | 0.285714286 | |
| X | 0 | 0 | 0 | | 0.090909091 | 0.09091 | 0 | 0.090909091 | |
| XI | 0 | 0 | 0 | | 0 | 0 | 0 | 0 | |
| XII | 0 | 0.0625 | 0.0625 | | 0.1 | 0.1 | 0 | 0.1 | |

| | WT (12/122/948) | | | | LacZ/LacZ (12/122/949) | | | | |
|---|---|---|---|---|---|---|---|---|---|
| | Tubules with gy | Tubules with sgl | Tubules with any | Total tubules | Tubules with gy | Total gy | Tubules with sgl | Tubules with any | Total tubules |
| I | 0 | 1 | 1 | 24 | 5 | 5 | 3 | 8 | 23 |
| II-III | 4 | 1 | 5 | 21 | 14 | 16 | 7 | 21 | 25 |
| IV | 7 | 2 | 9 | 16 | 6 | 8 | 6 | 12 | 12 |
| V-VI | 2 | 2 | 4 | 22 | 6 | 8 | 9 | 15 | 31 |
| VII | 0 | 1 | 1 | 13 | 1 | 11 | 2 | 3 | 13 |
| VIII | 0 | 0 | 0 | 13 | 1 | 6 | 0 | 1 | 8 |
| IX | 0 | 0 | 0 | 11 | 1 | 2 | 5 | 6 | 18 |
| X | 0 | 0 | 0 | 9 | 0 | 0 | 0 | 0 | 10 |
| XI | 0 | 0 | 0 | 10 | 0 | 0 | 0 | 0 | 11 |
| XII | 0 | 0 | 0 | 9 | 0 | 0 | 0 | 0 | 6 |
| | | | | 148 | | | | | 157 |
| | % Tubules with gy | % Tubules with sgl | % Tubules with any | Total % Tubules ( | % Tubules with gy (l | gy per tul | % Tubules with sgl (l | % Tubules with any (LacZ/LacZ) | Total % Tubules (LacZ/LacZ) |
| I | 0 | 0.041666667 | 0.041666667 | | 0.217391304 | 0.21739 | 0.130434783 | 0.347826087 | |
| II-III | 0.19047619 | 0.047619048 | 0.238095238 | | 0.56 | 0.64 | 0.28 | 0.84 | |
| IV | 0.4375 | 0.125 | 0.5625 | | 0.5 | 0.66667 | 0.5 | 1 | |
| V-VI | 0.090909091 | 0.090909091 | 0.181818182 | | 0.193548387 | 0.25806 | 0.290322581 | 0.483870968 | |
| VII | 0 | 0.076923077 | 0.076923077 | | 0.076923077 | 0.84615 | 0.153846154 | 0.230769231 | |
| VIII | 0 | 0 | 0 | | 0.125 | 0.75 | 0 | 0.125 | |

| | | | | | | | | |
|---|---|---|---|---|---|---|---|---|
| IX | 0 | 0 | 0 | | 0.055555556 | 0.11111 | 0.277777778 | 0.333333333 |
| X | 0 | 0 | 0 | | 0 | 0 | 0 | 0 |
| XI | 0 | 0 | 0 | | 0 | 0 | 0 | 0 |
| XII | 0 | 0 | 0 | | 0 | 0 | 0 | 0 |

| WT (12/122/952) | | | | | LacZ/LacZ (12/94/775) | | | | |
|---|---|---|---|---|---|---|---|---|---|
| | Tubules with gy | Tubules with sgl | Tubules with any | Total tubules | Tubules with gy | Total gy | Tubules with sgl | Tubules with any | Total tubules |
| I | 0 | 1 | 1 | 36 | 2 | 7 | 6 | 8 | 20 |
| II-III | 3 | 1 | 4 | 13 | 12 | 16 | 7 | 19 | 22 |
| IV | 5 | 2 | 7 | 17 | 3 | 9 | 5 | 8 | 8 |
| V-VI | 1 | 0 | 1 | 23 | 5 | 8 | 12 | 17 | 34 |
| VII | 0 | 0 | 0 | 12 | 3 | 3 | 5 | 8 | 27 |
| VIII | 0 | 0 | 0 | 13 | 3 | 3 | 1 | 4 | 16 |
| IX | 0 | 0 | 0 | 24 | 0 | 0 | 0 | 0 | 16 |
| X | 0 | 0 | 0 | 10 | 0 | 0 | 0 | 0 | 11 |
| XI | 0 | 0 | 0 | 10 | 0 | 0 | 0 | 0 | 8 |
| XII | 0 | 0 | 0 | 11 | 0 | 0 | 0 | 0 | 19 |
| | | | | **169** | | | | | **181** |
| | % Tubules with gy | % Tubules with sgl | % Tubules with any | Total % Tubules ( | % Tubules with gy ( | gy per tu | % Tubules with sgl ( | % Tubules with any (LacZ/LacZ) | Total % Tubules (LacZ/LacZ) |
| I | 0 | 0.027777778 | 0.027777778 | | 0.1 | 0.35 | 0.3 | 0.4 | |
| II-III | 0.230769231 | 0.076923077 | 0.307692308 | | 0.545454545 | 0.72727 | 0.318181818 | 0.863636364 | |
| IV | 0.294117647 | 0.117647059 | 0.411764706 | | 0.375 | 1.125 | 0.625 | 1 | |
| V-VI | 0.043478261 | 0 | 0.043478261 | | 0.147058824 | 0.23529 | 0.352941176 | 0.5 | |
| VII | 0 | 0 | 0 | | 0.111111111 | 0.11111 | 0.185185185 | 0.296296296 | |
| VIII | 0 | 0 | 0 | | 0.1875 | 0.1875 | 0.0625 | 0.25 | |
| IX | 0 | 0 | 0 | | 0 | 0 | 0 | 0 | |
| X | 0 | 0 | 0 | | 0 | 0 | 0 | 0 | |
| XI | 0 | 0 | 0 | | 0 | 0 | 0 | 0 | |
| XII | 0 | 0 | 0 | | 0 | 0 | 0 | 0 | |
